# South Western Ghats: a niche of clove (*Syzygium aromaticum* (L.) Merr. & Perry) diversification

**DOI:** 10.1101/2023.07.29.551085

**Authors:** G. S. Sreekala, M. Avinash, J. B. Reddappa, P. Reshma, J. Nainu, T. Anargha, J. S. Aswani

## Abstract

Survey conducted at major clove (*Syzygium aromaticum* (L.) Merr. & Perry) growing regions of South Western Ghats of India in the states of Kerala and Tamil Nadu identified thirty accessions with superior yield and distinct characters. Clove accessions were characterized based on twenty one qualitative and twelve quantitative traits. Variation among fifteen qualitative characters were found and the predominant traits observed were elliptical canopy shape, semi-erect branching pattern, lanceolate leaf lamina with acuminate apex, mid bud forming season, combination of 1,2,3,4,5 flower bud per cluster, medium sized bud, elliptical fruit and seed shape. Dendrogram constructed using the Unweighted Pair Group Method with Arithmetic Mean (UPGMA) method grouped thirty accessions into 5 major clusters at genetic similarity of 73%. Acc.19 was identified as a unique accession. Twelve quantitative characters were subjected to principal component analysis where three component groups were extracted, which explained 70.85 per cent of total variance. The score plot generated from the principal component loading grouped the accessions into 18 clusters. Minimal data set of four characters viz., plant height, canopy spread (EW), number of inflorescence per square meter and mature bud length was generated. Observation on the qualitative and minimum data set helps to identify the ideotypes. The geographical location was not found to influence genetic diversity.

**Highlight:**

- Clove genotypes in South Western Ghats were surveyed and diversity identified
- Combination of 1,2,3,4,5 flower bud per cluster was a unique character identified among cultivated clove
- Minimal data set of clove developed using plant height (m), canopy spread (EW), number of inflorescence per m^2^ and mature bud length (mm)
- Ideotype of clove developed

## INTRODUCTION

Clove (*Syzygium aromaticum* (L.) Merr. & Perry), native to the Moluccas islands of Indonesia (Alfian et al., 2019; Hariyadi et al., 2020; Mahulette et al., 2019a, 2021, 2022; Sundari et al., 2019), is a valuable spice crop that has been used as a food preservative and medicine for centuries. The unopened dried flower buds commonly used in the dishes impart a hot taste to the cuisines. Clove has high potential as a free radical scavenger due to its high antioxidant activity (Rojas et al., 2014). The world production of clove is estimated to be 1,81,788 MT (FAO 2022), with Indonesia being the leading producer. The domestic production of clove in India is 1,183 MT per annum (Spices Board 2022), and the cultivation is limited to parts of the Western Ghats of Tamil Nadu, Kerala, Karnataka, and the Andaman and Nicobar Islands.

In India, clove is cultivated in the hilly terrain of the Western Ghats at higher elevations in Tamil Nadu and Karnataka and in the red soils of the midlands of Kerala. The clove population in India originated from a few trees that were originally introduced. Moreover, the self pollinating nature of this crop limits the scope for variability and thus possesses a very narrow genetic base for clove in India. Plant characterization at the morphological level is a potential tool for the identification of clove diversity (Mahulette et al., 2022). The morphological diversity among various clove types might offer vital information needed for crop improvement (Alfian et al., 2019; Mahulette et al., 2019b; Yulianah et al., 2020; Mahulette et al., 2022).

Research works on the identification of clove accessions based on morphological markers are limited in India. Three distinctly different morphological variants in cloves were located during the surveys conducted in Tamil Nadu by Krishnamoorthy and Rema (1994) and the identified promising variants offered great potential for utilising their diversity for crop improvement programmes. Balakrishnamoorthy and Kennedy (1999) reported a few variants based on morphological and flower bud characteristics in a survey conducted in the Keeriparai and Maramalai regions of Kanyakumari in Tamil Nadu. In general, differences have been recorded in the shape of trees, bearing habits, cropping season, and variation in yield, color, shape and dimension of clove in India. The present study reports the morphological distinctiveness and genetic divergence based on qualitative and quantitative traits in the existing populations of clove accessions in the South Western Ghats of India, which is the major clove growing tract of India.

## MATERIALS AND METHODS

A survey was carried out in major clove growing areas of southern Kerala and Kanyakumari district of Tamil Nadu to identify the extent of genetic divergence in the existing populations of clove during 2017-18 (Figure 1).

**Figure 1.**
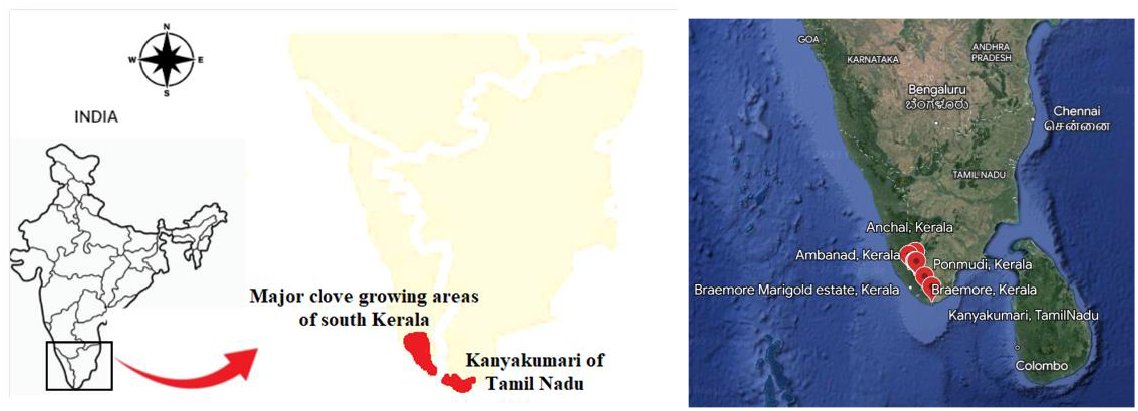
Study locations

From a survey of 1800 plants, thirty accessions that were superior in yield with distinct morphological characters were selected and characterized morphologically. The plant materials were selected from productive local trees that were more than 25 years old. All the selected accessions were subjected to morphological characterization based on tree, leaf, bud, flower, fruit and seed characters. Accessions were characterized with the help of descriptors of *Garcinia mangostana*, published by International Plant Genetic Resources Institute, Rome (IPGRI, 2003) and minimal descriptors for *Syzigium cumini* (PPV & FRA, 2012) and *Myristica fragrans* (PPV & FRA, 2012); Vikram (2016). The qualitative characters involved were canopy shape, branching pattern, color of young leaf, color of mature leaf, leaf lamina shape, leaf apex shape, leaf arrangement, bud forming season, bud clustering habit, bud size, position of flower, petal color, sepal color, color of stigma, color of peduncle, color of hypanthium, fruit shape, mature fruit color, ripe fruit color, seed shape and seed color. Canopy shape, branching pattern, leaf lamina and apex shape, leaf arrangement, bud forming season, bud clustering habit, position of flower, fruit shape and seed shape were recorded based on visual observation. The color of young leaf, mature leaf, petal, sepal, stigma, peduncle, hypanthium, fruit and seed were recorded as per the Royal Horticulture Society Color Charts. Bud size was measured during the harvesting period when they turned slightly pinkish color and were categorized as small, medium and large. The qualitative data were considered for cluster analysis using NTSYS (Numerical Taxonomy System) package 2.2. Observation on quantitative characters was measured and principal component analysis was carried out using KAU-GRAPES 1.0.0. Dry bud yield of the selected accessions was recorded for five consecutive years and the recorded data were subjected to pooled analysis.

## RESULTS AND DISCUSSION

### Morphological characterization of clove accessions

Considerable variation was present among the accessions for 15 out of 21 qualitative characters observed (Figure 2 to 4). However, uniformity occurred with leaf arrangement, position of flower, color of peduncle, mature fruit color, ripe fruit color and seed color among all the accessions (Table 1). The canopy shapes observed in the accessions were elliptical, cylindrical, conical and pyramidal with elliptical shapes being the most common (40%), followed by cylindrical, conical and pyramidal shapes. Canopy shapes associated with the branching pattern of a tree play a major role in the capture of solar radiation (Maiti et al., 2015). Majority of the clove accessions had semi erect branching pattern (56.67%), followed by irregular (33.33%) and erect (10%).

**Table 1.** Non variable characters observed among the selected clove accessions.

| Sl. No. | Characters | Expression |
| --- | --- | --- |
| 1 | Leaf arrangement | Opposite |
| 2 | Position of flower | Terminal |
| 3 | Colour of peduncle | Green |
| 4 | Mature fruit colour | Dark purple red |
| 5 | Ripe fruit colour | Bluish black |
| 6 | Seed colour | Green brown |

**Figure 2.**
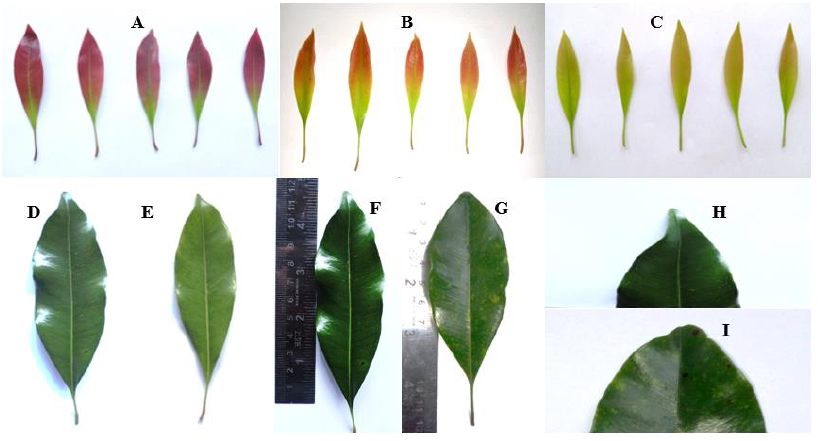
Expressions for leaf characters A. Red pink with light green tinged young leaves (Code: 43 A, B) B. Purple red with light green young leaves (Code: N 57A) C. Yellow green with light green young leaves (Code: 149 A) D. Dark green mature leaf (Code: 134 A) E. Green mature leaf (Code: 130 A) F. Lanceolate shape G. Narrowly Elliptic shape H. Acuminate tip I. Acute tip

**Figure 3.**
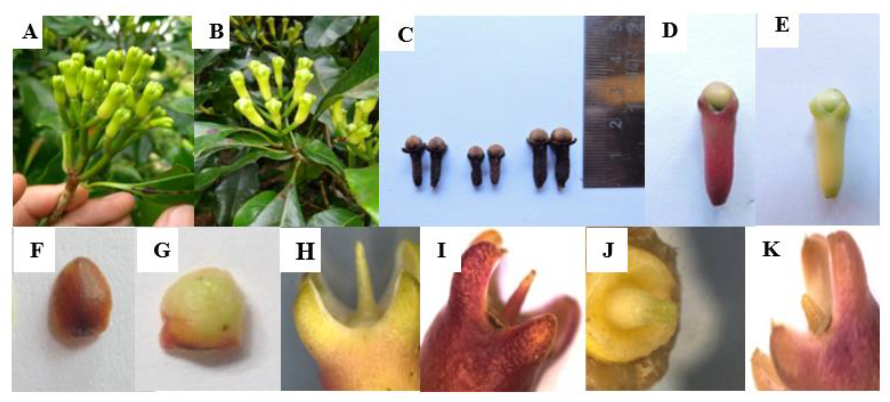
Expressions for floral characters A. Combination of 1,2,3,4,5 flower buds per cluster B. Combination of 1,2,3 flower buds per cluster C. Expression of bud size (small, medium & large) D. Purple red hypanthium E. Light red pink hypanthium F. Green brown (Code:152D) petal G. Light green Code:149D) petal H. Yellow green (Code:150C) sepal I. Dark purple red (Code:53B) sepal J. Light green (Code: 145A) K. Yellow green (Code: 154B)

Mature leaves of the trees were dark green (66.67%) or green (33.33%) while red pink with green tinge (90%), yellow green (6.67%), or purple red (3.33%) were observed among the young leaves. The predominant leaf character was lanceolate lamina with acuminate (86.67%) followed by narrowly elliptical with acute (13.33%) apex. The bud forming season in clove is categorized as early, mid and late. The early bud forming season in South Western Ghats started from September to October, mid season from November to December and late season from January to February. However, mid season bud forming was predominant (70%). A group of forest clove with large canopy was found to bloom early by Mahulette et al. (2019a).

Clove inflorescence is paniculate cymes and consists of many groups of three flower each. Bud clustering habit of the majority (86.67%) accessions observed were with combination of 1,2,3 flower buds per cluster and only 13.33% had a combination of 1,2,3,4,5 flower buds per cluster, which is a rare character. The majority of the accessions in our survey had medium sized buds. Balakrishnamoorthy and Kennedy (1999) reported three different types of clove flower buds *viz*., small, medium and big. The predominant hypanthium color was observed was light red pink. A unique accession, Acc. 19, exhibited dark purple red color. The majority had petal colour of light green and few green brown. Two types of sepal coloration were observed in the selected genotypes *viz*. yellow green (96.67%) and dark purple red (3.33%). Light green was the most common color of the stigma, followed by yellow green. Oblong shape of the fruit and seed were predominant and a few exhibited elliptic shape. Ravindran (2006) also reported oblong shaped seed in clove. The shape of the seed of clove was totally dependent on the fruit shape and oblong fruit produced oblong seeds and elliptical fruit produced elliptical seeds. Morphological variations among local clove accessions may due to genetic factors. But external environmental factors have a significant impact on the variation in phenotype among the genotypes (Adu et al., 2021; Ji et al., 2021; Luong et al., 2021; Tesfaye 2021; Mahulette et al., 2022).

### Diversity analysis of clove accessions

Jaccard’s similarity coefficient was calculated to determine the phylogenetic relationship and genetic divergence among the clove accession. The similarity coefficient ranged between 0.41 to 1.00 indicating that the clove accessions used in the present study had a low level of genetic variation. The maximum similarity value of 1.00 was observed between Acc. 5 and Acc. 6 and between Acc. 13, Acc. 17 and Acc. 25, whereas Acc.19 was quite distinct from other accessions, as indicated by the low value of the similarity coefficient (0.41). The Unweighted Pair Group Method with Arithmetic Mean (UPGMA) based dendrogram clustered the thirty accessions into five major clusters at 73% genetic similarity (Figure 5).

**Figure 4.**
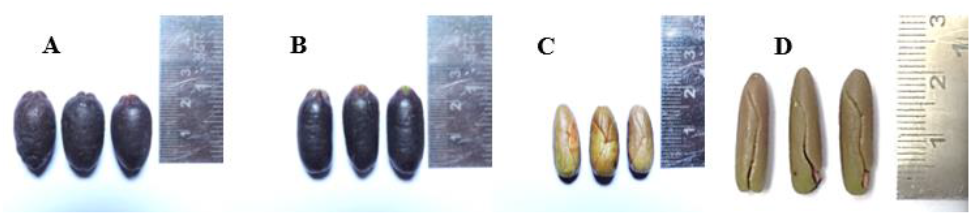
Expressions for fruit and seed characters A. Elliptical fruit B. Oblong fruit C. Elliptical seed D. Oblong seed

**Figure 5.**
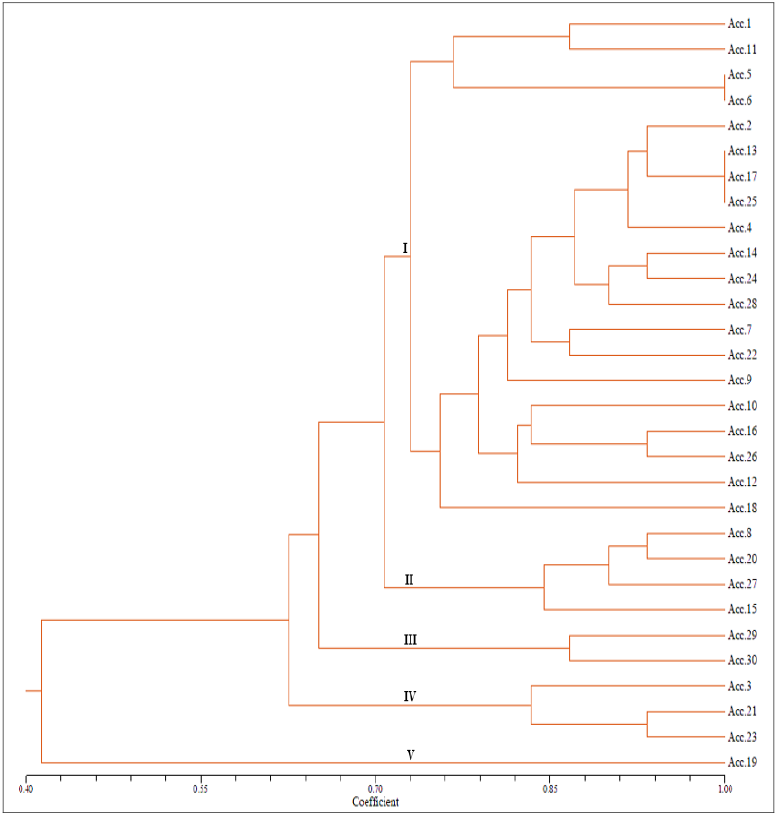
Dendrogram constructed by UPGMA clustering method based on qualitative morphological traits of thirty clove accessions

Cluster analysis grouping is more influenced by the number of similar characters (Alam et al., 2021; Jin et al., 2021; Kang et al., 2021; Liantoni et al., 2021; Loukili et al., 2021; Rathinavel, 2018; Zhang, 2021; Mahulette et al., 2022). The cluster I had 20 accessions, which indicates the absence of significant diversity for qualitative characters in clove. All the accessions possessed similarity among characters such as colour of young leaves (red pink with light green tinge), hypanthium (light red pink), petal (light green with pinkish tinge), sepal (yellow green), fruits (oblong) and seed (oblong).

Cluster II contained four accessions, Acc. 8, Acc. 20, Acc. 27, Acc. 15, which showed similar characters except canopy shape, branching pattern and mature leaf color. Cluster III contained two accessions, Acc. 29 and Acc. 30. These two accessions differed in only two characters; bud forming season and bud size. Acc. 29 was mid season bearer producing medium sized buds while Acc. 30 was an early bearer producing small buds. The cluster IV contained 3 accessions, Acc. 3, Acc. 21, Acc. 23, had all similar characteristics except canopy shape, branching pattern and bud forming season. Cluster V had only one accession, Acc. 19, identified as a unique accession having young leaf with purple red colour with light green tinge, dark purple red hypanthium, green brown petal, dark purple red sepal and yellow green stigma.

### Principle Component Analysis (PCA) of clove accessions

A multivariate analysis of twelve quantitative characters including dry bud yield of selected clove accessions was done (Table 2). Among the thirty accessions examined, length of mature bud ranged from 15.26mm to 19.06mm and the diameter varied from 4.90mm to 6.47mm. Analysis of the data on the pooled mean of dry bud yield recorded for five years selected clove accessions Acc. 1, Acc. 5, Acc. 6, Acc. 7 and Acc. 20 as high yielding ones (Table 3).

**Table 2.** Quantitative characterization of selected clove accessions.

| Accessions ID | Plant height (m) | Girth at 45 cm | Canopy spread |  | Number of branches | Number of inflorescence/ m<sup>2</sup> | Number of flower buds/ inflorescence | Single bud weight (fresh) (mg) | Single bud weight (dry) (mg) | Mature bud length (mm) | Mature bud diameter (mm) |
| --- | --- | --- | --- | --- | --- | --- | --- | --- | --- | --- | --- |
|  |  |  | NS | EW |  |  |  |  |  |  |  |
| Acc.1 | 9.20 | 118.5 | 4.20 | 3.75 | 48 | 114 | 13.47 | 327.05 | 114 | 19.06 | 5.94 |
| Acc.2 | 9.7 | 67.33 | 4.55 | 4.1 | 40 | 71 | 9.42 | 243.6 | 73 | 15.85 | 5.2 |
| Acc.3 | 13.6 | 138.1 | 4.60 | 4.35 | 44 | 73.75 | 8.47 | 400.37 | 128.5 | 17.73 | 6.41 |
| Acc.4 | 6.45 | 86.2 | 4.95 | 5.10 | 38 | 79.5 | 9.31 | 310.59 | 103 | 18.56 | 5.79 |
| Acc.5 | 6.55 | 85.4 | 5.15 | 4.30 | 46 | 70.25 | 15.05 | 367.47 | 101.5 | 18.76 | 5.83 |
| Acc.6 | 7.35 | 96 | 4.80 | 4.75 | 39 | 97.5 | 18.21 | 334.2 | 91.6 | 18.04 | 5.83 |
| Acc.7 | 14.95 | 99.8 | 7.42 | 7.05 | 47 | 43.5 | 10.47 | 324.48 | 110 | 18.73 | 5.55 |
| Acc.8 | 9.7 | 90.1 | 5.90 | 6.20 | 38 | 51 | 10.42 | 314.43 | 104 | 18.53 | 5.1 |
| Acc.9 | 10.81 | 114 | 6.1 | 5.9 | 46 | 75.25 | 6 | 328.4 | 94.8 | 17.27 | 6.04 |
| Acc.10 | 8.16 | 103.5 | 5.5 | 4.28 | 43 | 71.5 | 10.48 | 253.81 | 81.5 | 17.81 | 6.47 |
| Acc.11 | 9.83 | 122 | 4.06 | 3.77 | 39 | 86 | 14.52 | 276.25 | 83.4 | 16.73 | 5.49 |
| Acc.12 | 7.36 | 108.3 | 4.4 | 3.65 | 42 | 94.75 | 10.74 | 324.16 | 107.7 | 18.54 | 5.81 |
| Acc.13 | 12.1 | 134 | 6.1 | 7.52 | 48 | 73.25 | 10.62 | 358.52 | 108.8 | 18.70 | 5.84 |
| Acc.14 | 9.15 | 110.6 | 5.1 | 4.42 | 37 | 79 | 9.10 | 317.75 | 102.3 | 18.56 | 5.78 |
| Acc.15 | 7.49 | 73.2 | 4.9 | 4.63 | 32 | 74.75 | 9.44 | 354.22 | 98.5 | 17.93 | 5.53 |
| Acc.16 | 9.62 | 120.2 | 4.6 | 4.07 | 52 | 68.5 | 7.25 | 337.18 | 78.6 | 16.26 | 5.83 |
| Acc.17 | 8.7 | 102.4 | 5.4 | 4.51 | 45 | 73.25 | 11.81 | 274.3 | 112 | 18.70 | 5.54 |
| Acc.18 | 7.68 | 97.4 | 4.83 | 4.62 | 38 | 75 | 10.36 | 341.84 | 98.2 | 17.65 | 5.85 |
| Acc.19 | 5.6 | 53 | 3.25 | 2.90 | 31 | 48.5 | 7.47 | 237.04 | 76 | 15.99 | 4.9 |
| Acc.20 | 7.35 | 76.55 | 4.23 | 3.65 | 49 | 114.25 | 9.31 | 305.58 | 81 | 18.38 | 5.32 |
| Acc.21 | 6.85 | 102.5 | 6.15 | 7.00 | 41 | 87.75 | 7.1 | 358.3 | 99 | 18.03 | 5.2 |
| Acc.22 | 6.50 | 85.9 | 4.30 | 4.00 | 38 | 92.5 | 9 | 253.96 | 75.5 | 15.26 | 5.18 |
| Acc.23 | 5.15 | 68.2 | 3.45 | 3.55 | 41 | 99.5 | 9.57 | 337.59 | 89.5 | 18.18 | 5.63 |
| Acc.24 | 5.30 | 69 | 4.35 | 3.90 | 48 | 113 | 10.47 | 326.32 | 92.5 | 17.89 | 5.39 |
| Acc.25 | 7.85 | 82.9 | 5.05 | 5.8 | 43 | 79.5 | 12.1 | 332.35 | 94 | 17.97 | 5.9 |
| Acc.26 | 10.30 | 92.5 | 5.0 | 5.8 | 44 | 69 | 7.47 | 348.5 | 108 | 17.94 | 5.94 |
| Acc.27 | 5.9 | 65.8 | 4.25 | 4.05 | 36 | 100.5 | 12 | 330.5 | 94.5 | 18.77 | 5.27 |
| Acc.28 | 15.25 | 97.3 | 4.10 | 4.30 | 33 | 85 | 10.94 | 281 | 80 | 16.25 | 5.18 |
| Acc.29 | 7.15 | 55.3 | 3.15 | 3.40 | 31 | 24.75 | 6.68 | 274.53 | 82 | 15.68 | 5.19 |
| Acc.30 | 6.80 | 44.1 | 3.1 | 2.95 | 26 | 36.25 | 6.05 | 216.27 | 66.5 | 14.94 | 4.92 |

**Table 3.** Dry bud yield of selected clove accessions.

| Accessions ID | Dry bud yield per tree (kg / tree) |  |  |  |  |  |
| --- | --- | --- | --- | --- | --- | --- |
|  | 2016-17 | 2017-18 | 2018-19 | 2019-20 | 2020-21 | Pooled mean |
| Acc.1 | 7.3 | 6.7 | 7.5 | 6.54 | 5.1 | 6.63 |
| Acc.2 | 7.89 | 3.1 | 2.64 | 4.97 | 3.89 | 4.50 |
| Acc.3 | 7.12 | 3.1 | 3.32 | 5.98 | 3.98 | 4.7 |
| Acc.4 | 3.89 | 7.06 | 3.99 | 2.9 | 5.12 | 4.59 |
| Acc.5 | 5.46 | 6.12 | 8.04 | 5.45 | 5.97 | 6.21 |
| Acc.6 | 6.12 | 4.87 | 6.89 | 5 | 5.87 | 5.75 |
| Acc.7 | 8.98 | 3.1 | 7.3 | 6.7 | 3.1 | 5.84 |
| Acc.8 | 6.2 | 3.5 | 5 | 3.8 | 5.43 | 4.79 |
| Acc.9 | 6.93 | 3.2 | 4.9 | 4.2 | 3.0 | 4.45 |
| Acc.10 | 2.8 | 4.6 | 3.8 | 2.6 | 4.1 | 3.64 |
| Acc.11 | 4.7 | 3.6 | 4.9 | 6.8 | 4.03 | 4.81 |
| Acc.12 | 2.7 | 5.6 | 8.07 | 2.9 | 6.05 | 5.06 |
| Acc.13 | 3.2 | 6.01 | 3.5 | 3.8 | 6.5 | 4.60 |
| Acc.14 | 6.9 | 2.1 | 6.5 | 4.8 | 4.2 | 4.90 |
| Acc.15 | 3.2 | 3.7 | 5.02 | 2.1 | 6.23 | 4.05 |
| Acc.16 | 7.12 | 2.4 | 4.1 | 3.2 | 4.2 | 4.20 |
| Acc.17 | 3.9 | 7.3 | 3.01 | 2.6 | 5.1 | 5.28 |
| Acc.18 | 1.9 | 8.01 | 3.9 | 4.78 | 6.01 | 4.92 |
| Acc.19 | 3.02 | 1.9 | 2.9 | 2.1 | 3.3 | 2.64 |
| Acc.20 | 6.7 | 4.8 | 5.1 | 7.25 | 5.09 | 5.79 |
| Acc.21 | 3.0 | 5.5 | 5.9 | 3.1 | 4.2 | 4.34 |
| Acc.22 | 6.12 | 8.2 | 2.9 | 3.9 | 2.6 | 4.74 |
| Acc.23 | 3.5 | 5.3 | 3.6 | 4.3 | 3.9 | 4.12 |
| Acc.24 | 4.7 | 4.5 | 3.4 | 4.8 | 2.7 | 4.02 |
| Acc.25 | 4.5 | 3.2 | 3.2 | 3.5 | 2.4 | 3.36 |
| Acc.26 | 6.02 | 4.6 | 5.2 | 5.7 | 3.6 | 5.02 |
| Acc.27 | 6.89 | 5.46 | 2.6 | 4.5 | 3.7 | 5.02 |
| Acc.28 | 2.01 | 1.34 | 2.1 | 1.23 | 1.0 | 1.54 |
| Acc.29 | 3.8 | 2.6 | 2.5 | 2.9 | 5.8 | 3.52 |
| Acc.30 | 1.66 | 3.9 | 5.76 | 1.03 | 5.89 | 3.65 |

PCA was carried out for twelve quantitative characters which included plant height, girth at 45cm, canopy spread NS, canopy spread EW, number of branches, number of inflorescences per m^2^, variation in the number of flower buds per inflorescence, single bud fresh weight, single bud dry weight, mature bud length, mature bud diameter and dry bud yield (pooled mean). PCA method had been universally used as a data reduction tool. Three principal component groups were selected from the component loading of the twelve quantitative characters with the criteria of Eigen values more than 1 (Table 4). Eigen values indicate the amount of the variance explained by each factor. Factors with eigen values greater than one explained more overall variation in the data than factors with eigen values less than one. Therefore, only factors with eigen values greater than one were retained for interpretation. These three components could explain 70.85% of total variance. Based on the data from principal component loadings (Table 5), score plot and biplot were formed for better understanding of the results.

**Table 4.** Eigen values, percentage of variance and cumulative percentage of variance of Principal Components.

| Principal component | Eigen value | Percentage of variance | Cumulative percentage of variance |
| --- | --- | --- | --- |
| PC1 | 5.371 | 44.755 | 44.755 |
| PC2 | 2.041 | 17.01 | 61.765 |
| PC3 | 1.09 | 9.085 | 70.85 |
| PC4 | 0.861 | 7.172 | 78.022 |
| PC5 | 0.703 | 5.858 | 83.88 |
| PC6 | 0.626 | 5.216 | 89.096 |
| PC7 | 0.472 | 3.932 | 93.028 |
| PC8 | 0.295 | 2.462 | 95.489 |
| PC9 | 0.252 | 2.098 | 97.587 |
| PC10 | 0.145 | 1.211 | 98.799 |
| PC11 | 0.082 | 0.682 | 99.481 |
| PC12 | 0.062 | 0.519 | 100 |

**Table 5.** Component loadings and percentage contribution of each variable to three principal components.

| Variables | Characters | Component loadings |  |  | Percentage contribution of variables |  |  |
| --- | --- | --- | --- | --- | --- | --- | --- |
|  |  | PC1 | PC2 | PC3 | PC1 | PC2 | PC3 |
| X1 | Plant height (m) | -0.178 | **0.464** | **0.385** | 3.169 | **21.553** | **14.794** |
| X2 | Girth at 45 (cm) | **-0.332** | 0.136 | **0.47** | **11.047** | 1.842 | **22.071** |
| X3 | Canopy spread (NS) | **-0.318** | 0.314 | -0.272 | **10.14** | 9.849 | 7.402 |
| X4 | Canopy spread (EW) | -0.278 | **0.384** | -0.341 | 7.713 | **14.775** | 11.623 |
| X5 | Number of branches | -0.308 | -0.098 | 0.205 | 10.04 | 0.958 | 4.185 |
| X6 | Number of inflorescence (m²) | -0.147 | **-0.508** | 0.193 | 2.147 | **25.778** | 3.706 |
| X7 | Number of flower buds per inflorescence | -0.184 | -0.35 | 0.07 | 3.394 | 12.245 | 0.494 |
| X8 | Single bud weight fresh (mg) | **-0.334** | -0.052 | -0.109 | **11.18** | 0.275 | 1.198 |
| X9 | Single bud weight dry (mg) | **-0.358** | 0.033 | -0.161 | **12.789** | 0.107 | 2.586 |
| X10 | Mature bud length (mm) | **-0.348** | -0.2 | -0.336 | **12.123** | 4.001 | 11.292 |
| X11 | Mature bud diameter (mm) | -0.303 | -0.04 | **0.396** | 9.186 | 0.159 | **15.717** |
| X12 | dry bud yield (kg / tree) | -0.264 | -0.291 | -0.222 | 6.975 | 8.457 | 4.933 |

The highest percentage of variance (44.75) was noted in principal component group 1 (PC1. The first principal component was positively contributed mainly by single bud weight dry, mature bud length, single bud weight fresh, girth at 45 (cm) and canopy spread (NS). The principal component group 2 contributed 17.01% of variation with highest percentage contribution from number of inflorescence (m^2^) with 25.778%, plant height with 21.553% and canopy spread (EW) at 14.775%. The third principal component PC3 is positively associated with girth at 45cm, mature bud diameter and plant height. The relation between the thirty accessions was plotted in the score plot developed from principal component 1 and 2 which contributed 61.765% variability (Figure 6).

**Figure 6.**
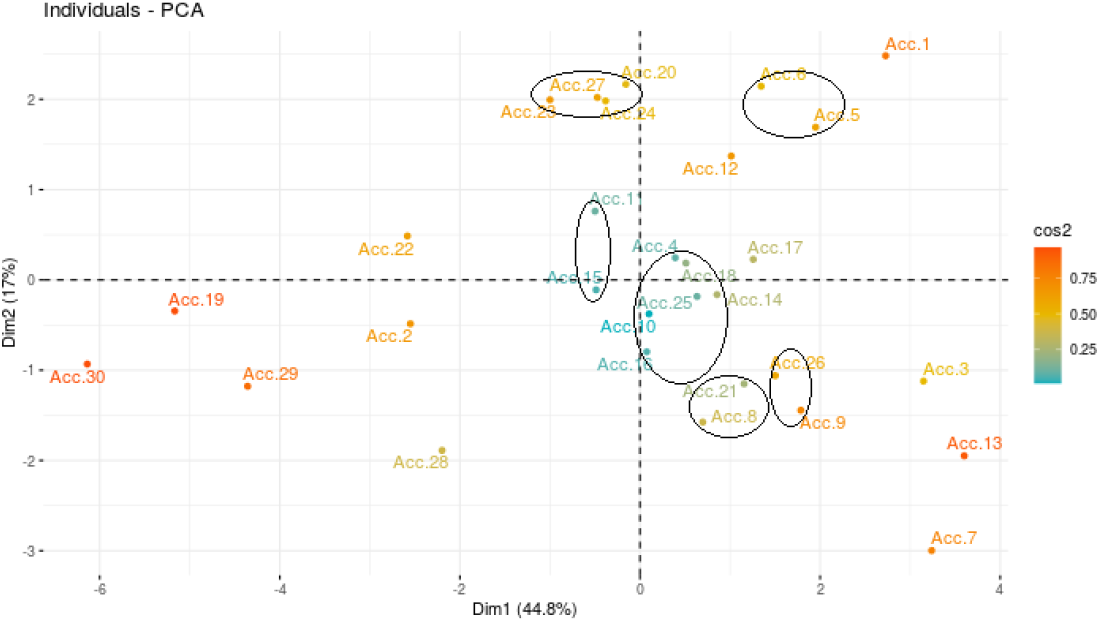
Clustering of accessions based on score plot

The function of a score plot is to group similar accessions and from the developed score plot, eighteen clusters were formed. Samples to the right in the score plot have high values for variables. The biplot generated from the principal component group 1 and 2 showed the characters that contribute more to variability (Figure 7). From the biplot, it was clear that number of inflorescence (m^2^), mature bud length, single bud weight fresh, single bud weight dry, girth at 45cm, canopy spread (NS), canopy spread (EW) and plant height contributed to more variability. The outcomes of the principal component analysis revealed the traits which contributed more to the variation.

**Figure 7.**
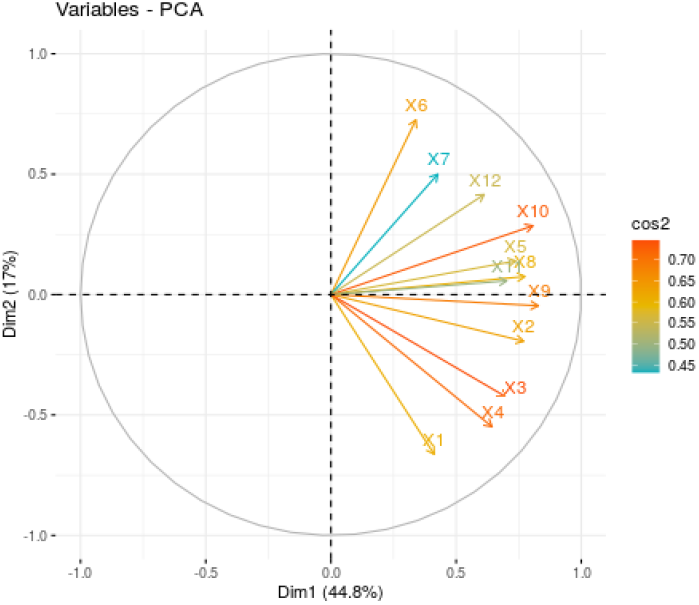
Biplot generated based on first two principal components

### Development of minimum data set (MDS) indicators for clove

The Minimum data set indicators were selected using the Principal Component Analysis (PCA)/correlation coefficient analysis approach from eight variables with the highest percentage contribution in the principal component (PC) groups PC1, PC2 and PC3. Only highly weighted factors within each principal component (PC) were retained for the development of the minimum data set (Mazumdar *et al*., 2014). The characters were X1-plant height, X2-girth at 45, X3-canopy spread (NS), X4-canopy spread (EW), X6-number of inflorescence, X8-single bud weight fresh, X9-single bud weight dry and X10-mature bud length. A correlation matrix for the twelve quantitative characters was worked out (Table 6). When the correlation value was less than 0.6, both the variables were selected whereas when the correlation value was more than 0.6, the highly weighed variables based on the percentage contribution of variables on principal components PC1, PC2 and PC3 were selected. From the eleven quantitative characters contributing to yield, four characters namely, X1-plant height, X4-canopy spread (EW), X6-number of inflorescences per m^2^ and X10-mature bud length were selected as a MDS for clove. The quantitative data of the showed that Acc. 1, Acc. 5, Acc. 6, Acc. 7 and Acc. 20 are the high yielding ones wherein Acc.1 had the highest value for single bud weight dry and mature bud length. Acc. 5 had the highest value for single bud fresh weight, Acc. 6 had the highest number of flower buds/ inflorescence, Acc. 7 had the highest value for canopy spread (NS) and Acc. 20 had the highest value for number of inflorescence per m^2^.

**Table 6.** Correlation matrix of the quantitative characters.

|  | X1 | X2 | X3 | X4 | X5 | X6 | X7 | X8 | X9 | X10 | X11 | X12 |
| --- | --- | --- | --- | --- | --- | --- | --- | --- | --- | --- | --- | --- |
| X1 | 1 | 0.602*** | 0.464** | 0.456* | 0.238 | -0.224 | 0.018 | 0.178 | 0.334 | 0.036 | 0.257 | -0.044 |
| X2 | 0.602**<br>* | 1 | 0.517** | 0.438* | 0.581*** | 0.23 | 0.267 | 0.494** | 0.582*** | 0.378* | 0.667*** | 0.318 |
| X3 | 0.464** | 0.517** | 1 | 0.881*** | 0.494** | -0.072 | 0.166 | 0.419* | 0.523** | 0.539** | 0.361* | 0.345 |
| X4 | 0.456* | 0.438* | 0.881*** | 1 | 0.354 | -0.117 | 0.05 | 0.481** | 0.488** | 0.441* | 0.228 | 0.165 |
| X5 | 0.238 | 0.581*** | 0.494** | 0.354 | 1 | 0.42* | 0.238 | 0.496** | 0.419* | 0.507** | 0.55** | 0.517** |
| X6 | -0.224 | 0.23 | -0.072 | -0.117 | 0.42* | 1 | 0.44* | 0.311 | 0.14 | 0.415* | 0.212 | 0.309 |
| X7 | 0.018 | 0.267 | 0.166 | 0.05 | 0.238 | 0.44* | 1 | 0.208 | 0.221 | 0.447* | 0.278 | 0.446* |
| X8 | 0.178 | 0.494** | 0.419* | 0.481** | 0.496** | 0.311 | 0.208 | 1 | 0.733*** | 0.653*** | 0.544** | 0.389* |
| X9 | 0.334 | 0.582*** | 0.523** | 0.488** | 0.419* | 0.14 | 0.221 | 0.733*** | 1 | 0.772*** | 0.546** | 0.508** |
| X10 | 0.036 | 0.378* | 0.539** | 0.441* | 0.507** | 0.415* | 0.447* | 0.653*** | 0.772*** | 1 | 0.469** | 0.585*** |
| X11 | 0.257 | 0.667*** | 0.361* | 0.228 | 0.55** | 0.212 | 0.278 | 0.544** | 0.546** | 0.469** | 1 | 0.308 |
| X12 | -0.044 | 0.318 | 0.345 | 0.165 | 0.517** | 0.309 | 0.446* | 0.389* | 0.508** | 0.585*** | 0.308 | 1 |

### Developing ideotype in clove accessions

An investigator shall first identify a clove tree that possesses a desirable set of qualitative characters and the quantitative characteristics which are mapped accordingly. In the bearing season the minimum data set characters shall be observed and recorded. The minimum data set characters that can taken into consideration would be plant height (m), canopy spread (EW), number of inflorescence per m^2^ and mature bud length (mm).

Thus by observing the qualitative characters and the minimum data set characters in the bearing season one can optimally sort out the ideotype clove accessions.

## FUNDING

Kerala State Planning Board has provided the financial assistance to conduct the study.

## Abbreviations

IPGRI: International Plant Genetic Resources Institute
NTSYS: Numerical Taxonomy System
UPGMA: Unweighted Pair Group Method with Arithmetic Mean
RAPD: Random Amplification Polymorphic DNA
PCA: principal component analysis
MDS: minimum data set

## Acknowledgments

We would like to thank the Kerala Agricultural University for providing the facilities for doing the research and the Kerala State Planning Board for funding this study through a research project. We also thank the clove estates situated in the south western ghats where the study was undertaken.

## Notes

### Competing Interest Statement

The authors have declared no competing interest.

